# Mapping DNA Methylation to Cardiac Pathologies Induced by Beta-Adrenergic Stimulation in a Large Panel of Mice

**DOI:** 10.1101/2024.10.25.619688

**Authors:** Caitlin Lahue, Eleanor Wong, Aryan Dalal, Wilson Tan Lek Wen, Shuxun Ren, Roger Foo, Yibin Wang, Christoph D Rau

## Abstract

**Background:** Heart failure (HF) is a leading cause of morbidity and mortality worldwide, with over 18 million deaths annually. Despite extensive research, genetic and environmental factors contributing to HF remain complex and poorly understood. Recent studies suggest that epigenetic modifications, such as DNA methylation, may play a crucial role in regulating HF-associated phenotypes. In this study, we leverage the Hybrid Mouse Diversity Panel (HMDP), a cohort of over 100 inbred mouse strains, to investigate the role of DNA methylation in HF progression.

**Objective:** We aim to identify epigenetic modifications associated with HF by integrating DNA methylation data with gene expression and phenotypic traits. Using isoproterenol (ISO)-induced cardiac hypertrophy and failure in HMDP mice, we explore the relationship between methylation patterns and HF susceptibility.

**Methods:** We performed reduced representational bisulfite sequencing (RRBS) to capture DNA methylation at single-nucleotide resolution in the left ventricles of 90 HMDP mouse strains under both control and ISO-treated conditions. We identified differentially methylated regions (DMRs) and performed an epigenome-wide association study (EWAS) using the MACAU algorithm. We identified likely candidate genes within each locus through integration of our results with previously reported sequence variation, gene expression, and HF-related phenotypes. *In vitro* approaches were employed to validate key findings, including gene knockdown experiments in neonatal rat ventricular myocytes (NRVMs). We also examined the effects of preventing DNA methyltransferase activity on HF progression.

**Results:** Our EWAS identified 56 CpG loci significantly associated with HF phenotypes, including 18 loci where baseline DNA methylation predicted post-ISO HF progression. Key candidate genes, such as Prkag2, Anks1, and Mospd3, were identified based on their epigenetic regulation and association with HF traits. In vitro follow-up on a number of genes confirmed that knockdown of Anks1 and Mospd3 in NRVMs resulted in significant alterations in cell size and blunting of ISO-induced hypertrophy, demonstrating their functional relevance in HF pathology.

Furthermore, treatment with the DNA methyltransferase inhibitor RG108 in ISO-treated BTBRT mice significantly reduced cardiac hypertrophy and preserved ejection fraction compared to mice only treated with ISO, highlighting the therapeutic potential of targeting DNA methylation in HF. Differential expression analysis revealed that RG108 treatment restored the expression of several methylation-sensitive genes, further supporting the role of epigenetic regulation in HF.

**Conclusion:** Our study demonstrates a clear interplay between DNA methylation, gene expression, and HF-associated phenotypes. We identified several novel epigenetic loci and candidate genes that contribute to HF progression, offering new insights into the molecular mechanisms of HF. These findings underscore the importance of epigenetic regulation in cardiac disease and suggest potential therapeutic strategies for modifying HF outcomes through targeting DNA methylation.

## Introduction

Heart failure (HF) is a leading cause of worldwide mortality and morbidity, associated with over 18 million deaths per year worldwide^1^. In the United States alone, approximately 6 million individuals are currently living with HF and HF is reported to play a role in approximately 1 in 8 deaths each year^1^. Heart Failure is a final unifying pathway for a number of distinct inciting etiologies and is typically diagnosed in the elderly after significant cardiac damage has already occurred^2^. This late age of detection results in a high degree of inter-individual environmental variation that impedes the efforts of scientists to identify genetic variants which underlie HF^2–4^. In earlier work, we used a large cohort of inbred mouse strains, the Hybrid Mouse Diversity Panel (HMDP) to circumvent these sources of environmental noise^5,6^. The HMDP consists of over 100 inbred strains of mice and contains approximately 4.2 million polymorphisms^7^. In our prior study, we used chronic beta adrenergic overdrive through the use of isoproterenol (ISO) to induce cardiac hypertrophy and failure in 104 strains of the HMDP. Through genetic mapping we identified 41 genome-wide significant loci in HF-associated phenotypes^5,6^ and, after combining our data with strain-and-condition-specific RNA transcriptomes, successfully identified and validated candidate genes at these loci through a combination of *in vitro* and *in vivo* approaches.

Recent research into HF has extended into a study of the epigenome, looking for non-sequence-level variations in DNA that are linked to changes in HF-associated phenotypes^8,9^. The DNA methylome, notably methylation of cytosines in CG dinucleotide pairs (CpGs), has been demonstrated to play a key role in the development of the heart and regulation of HF^8,10–13^, and epigenome-wide association studies (EWAS) have successfully identified specific CpGs linked to phenotypic changes during HF progression^13^. In past work we demonstrated that methylome differences between the inbred mouse strains BUB/J and Balb/cJ could be linked to ISO-induced HF susceptibility^14^.

In this study, we integrate DNA methylation captured at single nucleotide resolution from the left ventricles of control and ISO-treated hearts across 90 strains of the HMDP with gene expression and phenotypic traits from these strains and uncover convincing patterns of differentially methylated regions (DMRs) that correspond with disease severity. Application of the EWAS algorithm MACAU^15^ identified 56 CpG loci that are significantly associated with HF phenotypes, including 18 that link pre/un-treated DNA methylation status to post-ISO HF progression and severity. Through the use of a prioritization algorithm that links sequence variation, CpG methylation, gene expression, and phenotypic traits, we identify many high confidence EWAS candidate genes, including *Prkag2*, *Anks1*, and *Mospd3*. Using *in vitro* and *in silico* approaches, we validate the role of several of these genes in HF. Finally, we demonstrate that blocking the action of methyltransferases is sufficient to prevent cardiac hypertrophy in a murine strain (BTBRT<+>tf/J) that otherwise responds strongly to catecholamine overdrive. Our findings clearly demonstrate an interplay between DNA methylation, gene expression, and HF-associated phenotypes and represent a rich resource for future scientific study.

## Methods

### Hybrid Mouse Diversity Panel Isoproterenol Study

We previously reported^5,6,16^ a genetic study of heart failure in the Hybrid Mouse Diversity Panel, in which 8-10 week old (average 9.1 weeks) female mice from 105 diverse inbred mouse strains were divided into control (2 mice) and treated (4 mice) groups per strain. Treated mice were administered the β-adrenergic agonist isoproterenol (ISO) via intraperitoneally-implanted osmotic micropumps (Alzet, model 2004) at a rate of 30 mg ISO/ kg body weight/ day for 21 days, at which point all mice were sacrificed, organs removed, weighed, and flash frozen in liquid nitrogen. All mice were obtained from Jackson Labs or directly from the UCLA HMDP colony as described^5^. All mice were maintained on a standard chow diet and housed under pathogen-free conditions according to NIH guidelines. Mice underwent echocardiography before surgery and weekly thereafter until sacrifice at 21 days. Sections from the left ventricle of the heart were studied using Masson-Trichrome staining to quantify fibrosis levels as previously described^16^.

### RG108 Mouse Models

BTBRT<+>/tfJ and C57BL/6J female mice aged 8-10 weeks were obtained from Jackson Laboratories. The use of the mice were under the care and guidelines of National University of Singapore Institutional Animal Care and Use committee (NUS IACUC). 2mg of non-specific DNMT inhibitor *N*-phthalyl-_L_-trytophan (RG108) (Vector Biomed) was dissolved in 33 µl dimethyl sulfoxide (DMSO) (Sigma-Aldrich) and 15 µl ethanol^17^. For every 100 µl of RG108 mixture, 840 µl corn oil (Sigma-Aldrich) was added. The animals were divided into 3 groups (n=12 per group). Group 1 received saline to serve as baseline control. In group 2, the mice were implanted with an Alzet osmotic pump (model 2004) to deliver a consistent dose of ISO at 30mg/kg/day for 3 weeks. In group 3, the mice received ISO (30mg/kg/day) and a single (daily) subcutaneous dose of RG108 (12.5mg/Kg/day) for 3 weeks. Echocardiography was performed on all the animals at a weekly interval and sacrificed at week 3 post-implantation. The hearts were removed and weighed and the left ventricles (LV) were harvested for histology staining. DNA and RNA were isolated for RRBS-seq and RNA-seq assays.

### DNA Isolation and Reduced Representational Bisulfite Sequencing

DNA from 90 HMDP strains (see Supplemental Table 1) as well as from the RG108 cohorts were isolated from control and ISO-treated left ventricles. Each sample was lysed in RLT Buffer using a roto-stator homogenizer and processed using dnEasy kits (Qiagen) according to manufacturer instructions approach and quantified using the Qubit dsDNA HS Kit. RRBS-seq was performed as described in Gu et al^18^ with modifications. Briefly, 50 ng of purified DNA was digested with MspI (Fast digest MspI, Thermo Fisher Scientific FD0544) for 30 min at 37°C followed by heat inactivation at 65°C for 5 min. Lambda DNA (Thermo Fisher Scientific) was spiked into the DNA sample to serve as an internal control to calculate the bisulfite conversion efficiency. Library preparation was performed using NEBNext Ultra DNA library prep kit for Illumina (New England BioLabs) and ligated with methylated adapters for Illumina sequencing at a dilution of 1:10 (New England BioLabs). The adapter ligated DNA was subjected to bisulfite conversion with EpiTect fast bisulfite conversion kit (Qiagen) using the following cycling conditions: 2 cycles of (95°C; 5min, 60°C; 10min, 95°C; 5min, 60°C; 10min) and hold at 20°C. Bisulfite converted DNA was PCR amplified for 14-16 cycles using 2.5 U of Pfu Turbo Cx Hotstart DNA polymerase (Agilent Technologies, 600410) and size selected for fragments between 200 bp to 500 bp with Ampure Xp magnetic beads (Agencourt). Purified DNA was subjected to single end sequencing using the Illumina Hiseq 2500 at 1x 101 bp read length.

### RNA-seq Library Preparation and Data Analysis

RNA was isolated from the left ventricle of RG108 cohort animals using rnEasy kits (Qiagen). RNA-seq was performed with 1µg of total RNA using the Illumina Truseq kit according to manufacturer’s protocol. The library was subjected to paired-end sequencing on the Illumina Hiseq2500 at 2x 101 bp read length. RNA-Seq libraries were aligned to the mouse reference genome, mm10, using Tophat2 (version 2.2.0.12)^19^ with default parameters. The quality of the mapping was assessed using RNASEQC^20^. Gene expressions were computed using Cufflinks2 (version 2.2.1)^19^. Gene expression level was reported in Fragment Per Kilobase per Million Reads (FPKM).

### DNA Methylation Data Processing

RRBSseq reads were aligned to the mouse reference genome, mm10, using the BSseeker2 algorithm^21^ with default parameters. To ensure high data quality, CpGs with Q<30 and read depth of less than 3x were filtered out as well as CpGs in strains which had a detected SNP at the CpG site. Batch effects in the data were identified and corrected using COMBATseq^22^ (see Supplemental Figure 1A and 1B). To achieve an accurate estimate of methylation level, high read cutoffs were applied to eliminate PCR effects. CpGs having higher coverage than 99.9% percentile of other read counts were removed, using filterByCoverage function in methylKit^23^ package. Because methylation occurs almost exclusively in the CpG context, we focused only on cytosines in CpG dinucleotides (CGs).

**Figure 1.**
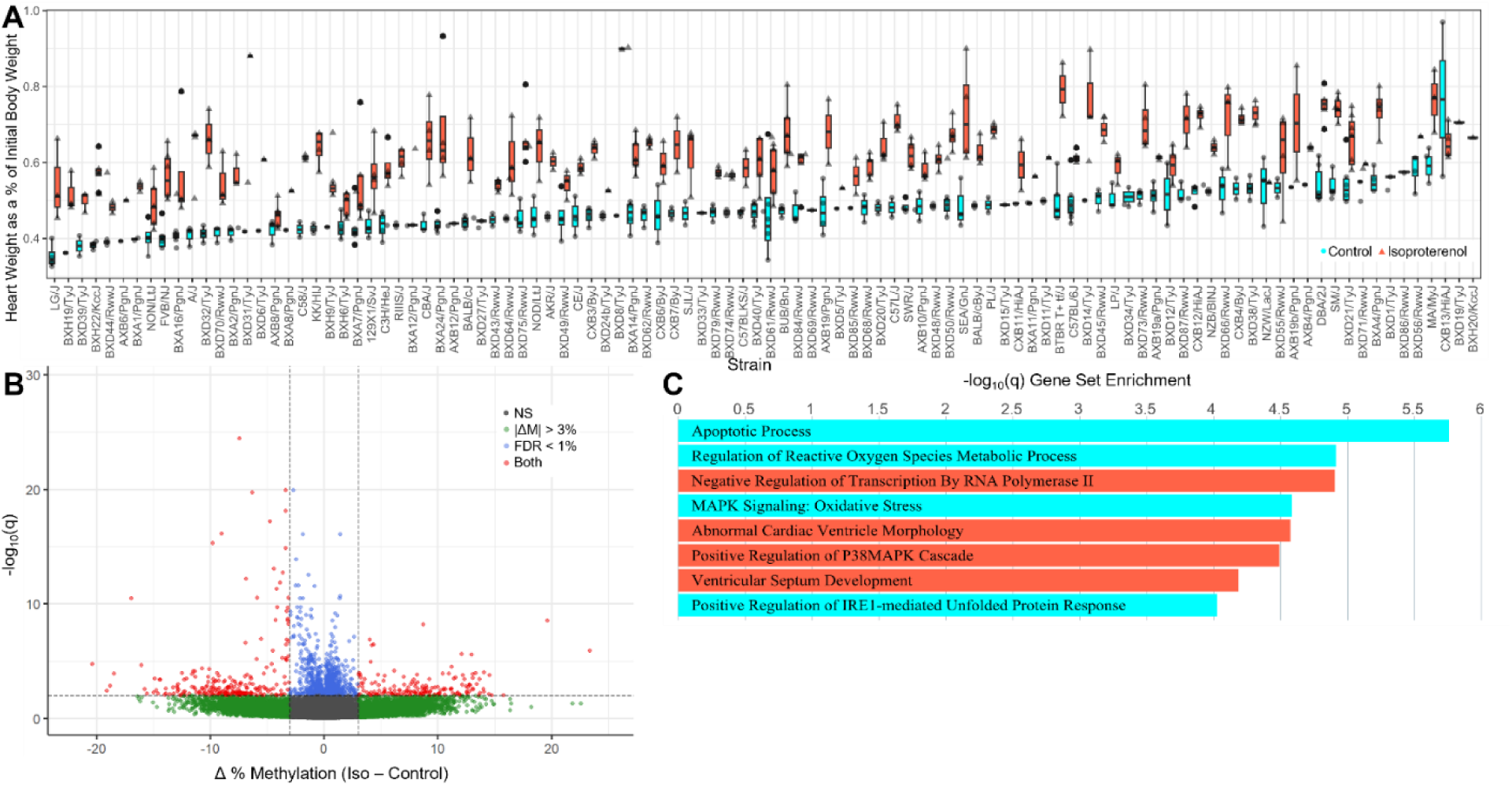
DNA Methylation Changes from ISO in the HMDP. **A)** Total Heart Weight as a percentage of day 0 body weight across 90 strains of the HMDP. **B)** Volcano plot showing differential methylation of CpGs with and without ISO. Green points are CpGs whose methylation shifts by at least 3% between conditions, while blue are CpGs that pass a 1% FDR threshold and red points are the 397 CpGs that meet both criteria. **C)** Gene Set Enrichment of genes proximal to significantly differentially methylated CpGs, with blue representing sites that are hypermethylated while red reflects sites that are hypomethylated.

### Identification of Differentially Methylated Regions

Percent methylation (PM) was calculated for each covered C by taking the ratio of methylated Cs divided by the total number of reads at that location. We then further limited our study to regions with at least 5x CpG coverage detected in 70% or more of the HMDP strains and used the ggbiplot^24^ R package to remove obvious outliers.

#### Differential methylation between Control and Isoproterenol-treated hearts

For each remaining CpG site in the dataset, we calculated differential methylation with the Methylkit R package^23^, which uses a logistic model to ascertain whether or not ISO has had an effect on methylation levels by modeling the log odds ratio based on the methylation proportion of a CpG *π*_*i*_ with or without the addition of a treatment term, or in other words whether 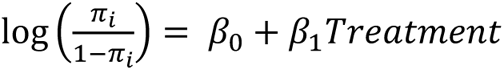 is a better model than 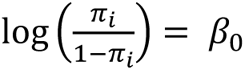. We considered all sites with a minimal shift of methylation of 3% and FDR < 1% for further study.

#### Differential methylation across the HDMP cohort

We relied on hypervariability, a previously described measure of DNA methylation variability used in past methylation studies of the HMDP^25,26^ to identify CpGs for further study. Briefly, hypervariable sites are CpGs in which the percent methylation shifts by over 25% in at least 5% of the affected strains. We modeled this off of the standard use of a minor allele frequency cutoff of 5%, as we have used in prior SNP-based studies^5,6,27^.

### Epigenome Wide Association Studies

We used the methylation-specific binomial mixed model package MACAU^15^ to test for association and account for population structure and relatedness between the mouse strains. MACAU models each CpG as

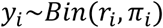

Where *r*_*i*_ is the total read count for the ith individual, *y*_*i*_ is the methylated read count for that individual, constrained to be an integer equal to or smaller than *r*_*i*_, and *π*_*i*_ is an unknown parameter that represents the true proportion of methylated reads for the individual at that site. MACAU then uses a logit link to model *π*_*i*_ as a linear function of parameters

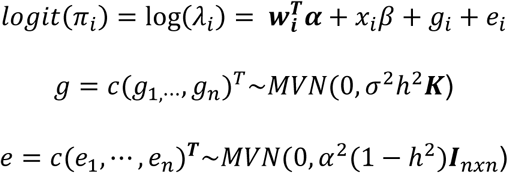

Where ***w***_***i***_ is a c-vector of covariates including an intercept and **α** is a c-vector of corresponding coefficients, *x*_*i*_ is the predictor of interest and β is its coefficient. g is a n-vector of genetic random effects that model correlation due to population structure and e is a n-vector of environmental residual errors that model independent variation. **K** is a known n x n relatedness matrix based, in our case, on genotype data, and standardized to ensure that *tr*(*K*)/*n* = 1 (this ensures that ℎ^2^ lies between 0 and 1 and can be interpreted as heritability). **I** is a n x n identity matrix, *σ*^2^ℎ^2^ is the genetic variance component, *σ*^2^(1 − ℎ^2^) is the environmental variance component, ℎ^2^ is the heritiability of the logit transformed methylation proportion (aka *logit*(*π*)) and MVN denotes the multivariate normal distribution.

To test for association of a CpG to a trait, MACAU tests the null hypothesis H0 : β= 0 for each site. It samples to compute an approximate maximum likelihood estimate *β̑*, its standard error *se*(*β̑*), and a corresponding p-value of significance as described^15^. Significant loci were determined by first calculating a Bonferroni-corrected significance threshold by dividing our alpha of 0.05 by the estimated number of correlated units of methylation, calculated as 3,330 (approximately 1 per 750 kb) in prior EWAS work in the HMDP^25^, resulting in a per-phenotype significance threshold of 1.5E-5. Although we have measured a total of 69 phenotypes, many of these are not independent, either linked to one another through physiology (e.g. LVID at diastole vs systole) or at times directly derived from combinations of other phenotypes (e.g. fractional shortening vs LVID). Through principle component analysis using the ggbiplot R package^24^, we estimated that we had approximately 36 “independent” phenotypes across our entire study. Therefore, to calculate our final threshold we performed another Bonferroni correction on our per-phenotype threshold to obtain a final threshold of 4.17E-7.

### Candidate Gene Selection

Previous reporting on DNA methylation in the HDMP^25^ identified the average correlation block size of a methylation locus (equivalent to the Linkage Disequilibrium block of a SNP locus) to be approximately 750kb{Orozco, 2015 #29}. To account for potentially larger loci, we extended our analysis to examine all genes that lay within 1 MB in either direction of the peak associated CpG in the locus, then leveraged other data from previously published HMDP cohorts to prioritize candidates^5,6^. First, we looked for mutations present in or around the promoter or exons of each gene that were predicted to cause a change in gene expression or function via the Wellcome Trust Mouse Genomes Resource^28,29^ which has fully sequenced each of the founder lines of the HMDP. Next, we looked whether the gene’s expression was significantly associated with the locus via eQTL^5^ or emQTL analyses. Next we examined whether the gene’s expression correlated with the phenotype associated with the locus^5,6^, and finally, whether there was literature evidence for association of this gene with the phenotype of interest or with DNA methylation. Genes with multiple lines of evidence, or strong evidence (e.g. very strong associations of gene expression with the locus) were prioritized for *in vitro* validation.

### *In vitro* Validation Studies

Neonatal Rat Ventricular Cardiomyocytes (NRVMs) were isolated from 1-4 day old rat neonates using the Cellutron Neomyocyte isolation kit (Cellutron) with modifications. Briefly, hearts were quickly removed and trimmed from neonatal rats and placed in ice cold PBS until 10 hearts had been isolated. PBS was removed and then replaced with 4mL digestion buffer, then incubated for 12 minutes at 37C on a stir plate at 150rpm in a 25 mL beaker with a 1’’ stir bar. This size beaker and stir bar was crucial for isolating large numbers of NRVMs. Supernatant was transferred to a new 15 mL tube and spun at 2,200rpm for 2 minutes. Supernatant was discarded and cells resuspended in digestion stop buffer with cell media at room temperature. Meanwhile, 4 mL of digestion buffer was added to the hearts and the entire process repeated 7-9 times until the heart turned a pale whitish-pink and fewer cells were recovered after centrifugation. All cells were centrifuge at 2,200 rpm for 2 minutes and then resuspended in 2 mL ADS buffer(12mM NaCl, 2mM HEPES, 1 mM NaH2PO4, 0.5mM Glucose, 0.5mM KCl, 0.1 mM MgSO4).

NRVMs were purified by passing them through a Percoll gradient, which was established by carefully layering 6mL of 1.059g/mL Percoll atop 3 mL of 1.082g/mL Percoll, both diluted in ADS buffer in a 15 mL conical tube. Cell suspension was slowly added to not disturb the layers, then centrifuged at 3000 rpm for 30 minutes at the slowest possible ramp up speed and with the brake disabled. Two bands of cells were visible, with cardiomyocytes concentrated in the lower band. Other cells were aspirated off and the NRVMs carefully extracted and diluted in 10 mL of ADS buffer followed by centrifugation at 2200 rpm for 3 minutes and supernatant discarded. y. NRVM pellet was then resuspended in 2 mL of DMEM with 10% FBS and 1% pen/strep and counted using a Countess II cell counter (ThermoFisher). Cells were plated onto gelatin-coated 12-well plates at a density of 200-250k cells per well.

We followed our previously established protocol for testing gene siRNAs in NRVMs (see Supplemental Table 2 for siRNAs). 24 hours after plating, DMEM media containing FBS and pen/strep was aspirated and wells washed 2x in PBS. DMEM media containing 1% ITS supplement (SigmaAldrich) was added to each well. That same day, siRNAs were transfected into cells using lipofectamine RNAiMax (Invitrogen) per manufacturer instructions. For each siRNA experiment, 6 wells across 2 12-well plates each got either control (no siRNA), scramble siRNA, or a siRNA obtained from IDT (See Supplemental Table 2). Transfections were allowed to proceed for 24 hours, then the media was refreshed and isoproterenol added to half of the wells at a final concentration of 60 mM. After 48 hours, photographs of each well were taken at 20x magnification and RNA isolated for qPCR validation of gene knockdown (see Supplemental Table S3). Cell cross-sectional area and confluence were assessed for each well by trained users.

### Gene Ontology Enrichment

Gene ontology enrichment was performed using the Gene Analytics Suite^30^ which uses a binomial test to test the null hypothesis that a defined set of genes is not over-represented within a given pathway and then corrected using the Benjamini-Hochberg correction (FDR). GeneAnalytics has several modules (e.g. a Gene Pathways module, a GO Terms module, etc.) We specify which module we use in the text as needed. All p values reported are corrected p values.

## Results

### HMDP Data Acquisition

We performed reduced representational bisulfite sequencing (RRBS) on 90 inbred mouse strains from the Hybrid Mouse Diversity Panel (HMDP) which we had previously used in a systems genetics study of beta-adrenergic driven cardiac hypertrophy and failure^5,6^ (See Supplemental Table 1 for mouse strains used in this study). 8-10 week old mice were divided into control and ISO-treated cohorts (30 mg/kg body weight/day via Alzet osmotic pumps).

Three weeks later, mice were sacrificed and isolated left ventricular DNA was cut with *MspI*, bisulfite converted, and size selected, followed by library prep and sequencing on an Illumina HiSeq2500 resulting in 174 100-bp single end libraries averaging 70.2 million reads per sample. Data was aligned to the mouse genome (mm10) using BSSeeker2^21^ with an average of 41.3 aligned reads across 2.8 million CpGs for an average mappability of 58.8% and an average coverage of 38x. We filtered out all (7,230) CpGs which had a polymorphism in the HMDP as detected by BSSeeker2. We then corrected for batch effects using COMBATseq^22^. We then limited our analysis to CpGs that were detected in at least 70% of the strains at 5x coverage, leaving us with a final number of 1.8 million CpGs for downstream analysis. The mouse genome consists of approximately 21.3 million CpGs, therefore we observed approximately 8.4% of all CpGs using RRBS.

Global methylation levels at CpGs shifted by -0.07% (standard deviation 1.8%, Figure 1B) in response to ISO challenge, suggesting that, at least globally, DNA methylation is not significantly affected by ISO. We identified a set of 168,251 hypervariable CpGs (>25% absolute change in variation in at least 5% (9) samples) for use in EWAS and other analyses.

For the same mouse strains, we measured 69 clinical traits, including heart and other organ weights, echocardiographic measurements and cardiac fibrosis, as well as gene expression using Illumina Mouse Ref 8.0 RNA microarrays as previously reported in our prior work on this cohort^5,6^ (see, for example, the variation observed in heart weights across the panel in Figure 1A).

### Observed Methylation Patterns Across the HMDP

Next, we examined the effects of ISO on our animals from a global perspective. (Figure 1B) We calculated the average methylation shift for each CpG between control and treated animals as well as the significance of this shift. We observe 27,603 CpGs that are nominally significant at p<0.05, and 1,413 CpGs which remain significant at an FDR of 1%. Overlapping these significant CpGs with the 18,723 CpGs which show an average shift of at least 3% between ISO and Control samples, we find 231 CpGs which are globally hypomethylated in response to ISO treatment and 166 CpGs which are globally hypermethylated in response to ISO at an FDR of 1% (Figure 1B). The nearest gene to each CpG was annotated using the genotate R package^31^ and gene ontology enrichments calculated using the GeneAnalytics platform’s Gene Ontology module^30^. Hypermethylated genes were enriched for, among other terms, Apoptosis (P=1.8E-6), oxidative stress (P=1.2E-5) and the unfolded protein response (P= 9.5E-5), while hypomethylated genes were enriched for RNA transcription (P=1.3E-5), Abnormal cardiac morphology (P=2.7E-5), and the P38MAPK cascade (P=3.2E-5) (Figure 1C, See Supplemental Table 4 for complete details).

Wanting a better sense of how genetic and environmental effects affected DNA methylation across the HMDP, we extracted the top 1% (1,623) hypervariable CpGs which showed the largest standard deviation across the HMDP (not necessarily between the Iso and Control cohorts) (Figure 2). Echoing what we observed when specifically focusing on CpGs which were affected by ISO treatment, we observe far stronger *strain* (genetic) effects on the methylome than *environmental* effects as evidenced by hierarchical clustering of the strain methylomes largely separating by genetic cohort rather than experimental condition. That is, we observe that each of the three Recombinant Inbred panels that make up the majority of the HMDP clustered independently, while the other inbred lines and the C57-associated lines formed their own clades in the strain dendrogram (Figure 2, bottom edge). By contrast, no clustering could be detected for isoproterenol status, with isoproterenol and control treated mice from the same strain tending to cluster together rather than separately (Figure 2 top edge). Among these top 1% varying CpGs, we observe 10 clusters across the HMDP panel. For each CpG, we identified the closest gene (if any) within 500kb of the CpG and then submitted these gene lists to the Pathway module of the Gene Analytics enrichment suite^30^ (Figure 2, right edge). Each cluster of CpGs we observed were enriched for one or more pathways, many of which are crucial to cardiac function. For example, we observe clusters involved in β adrenergic signaling (P=1.4E-6), Collagen Production (P=7.6E-5) and other major signaling or cytoskeleton-associated pathways (Full details in Supplemental Table 5).

**Figure 2.**
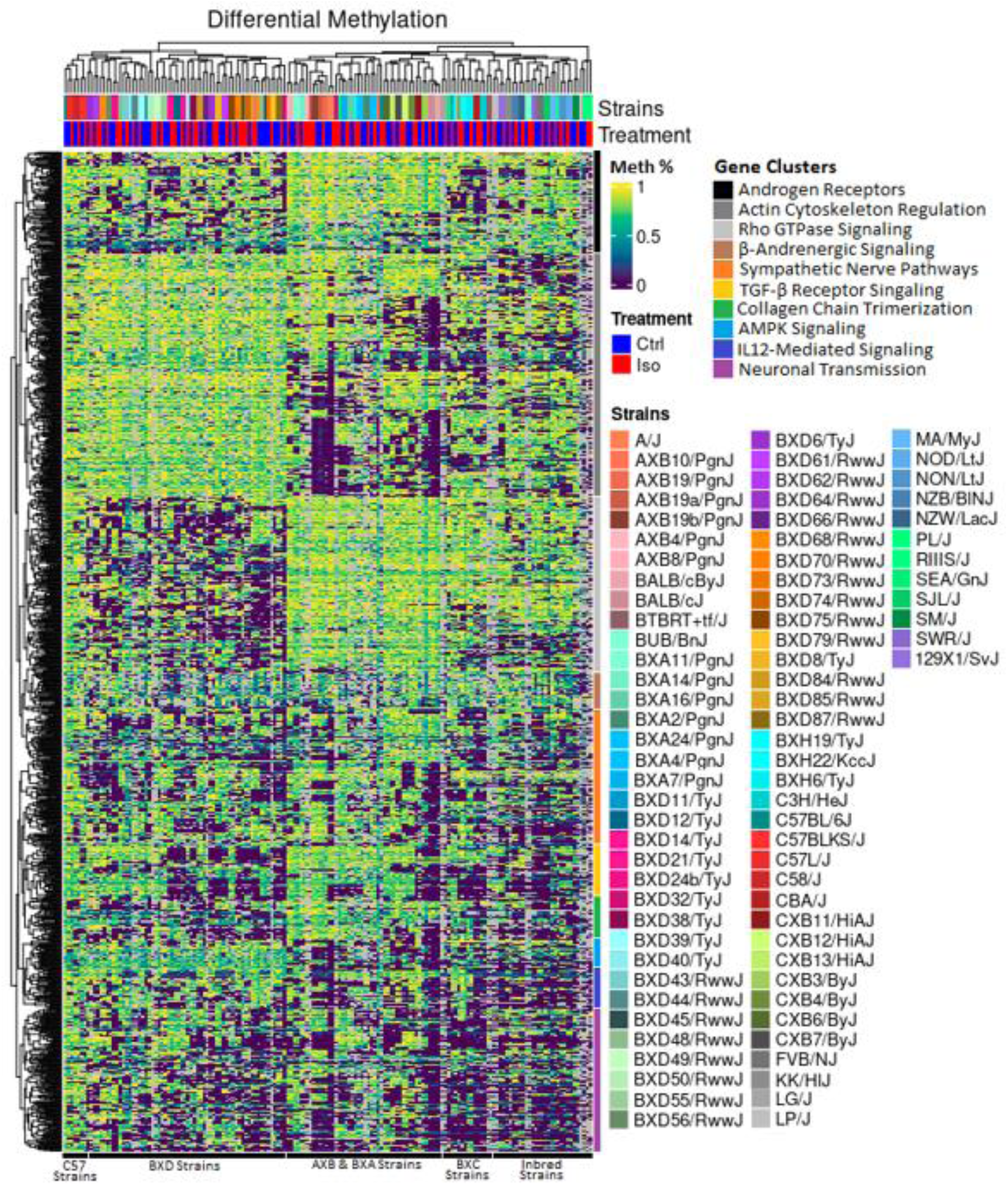
Heatmap of the top 1% of Differentially Methylated Loci across the HMDP Cohort. X axis is organized by hierarchical clustering and shows the separation of the panel into distinct cohorts based on their sub-panel origin, with the BXD, AXB/BXA and BxC RI panels clustering together (see bottom edge) while ISO treatment was not a major driver of clustering (top edge). CpGs were clustered along the Y axis based on similarity and genes proximal to these CpGs were analyzed for GO enrichments, seen along the right edge of the heatmap and annotated in the legend. Larger version available as Supplemental Figure 2.

### Variation in CpG Methylation is Associated with and Predictive of Heart-Failure-Associated Phenotypes

In order to identify associations between natural variation in CpG methylation across the HMDP and complex clinical traits, we performed a set of EWAS studies between Hypervariable CpG methylation and 69 traits, including heart and chamber weights, other organ weights, cardiac fibrosis, and echocardiographic parameters in control and ISO-treated animals as well as the change between ISO and control conditions (23 phenotypes each, see Supplemental Table 6). In contrast to our work with SNP-based GWAS in the HMDP, we elected to use a binomial mixed-model approach, MACAU, which was specifically designed for unsupervised determination of associations between CpGs and traits in WGBS and RRBS contexts. In keeping with best practices, we used a kinship matrix based on CpG methylation in contrast to a SNP-based kinship matrix^25^, which not only corrects for false associations caused by populations structure^7,27,32,33^ but also partially accounts for changes in tissue heterogeneity found in our RRBS data^34^.

Prior work in the realized HMDP has identified an average correlation structure in CpG methylation data (roughly equivalent to the concept of ‘linkage disequilibrium’ in SNP data) of 750kb^25^. We used this as a basis for determining a significance threshold in our data, performing Bonferroni-corrections on our initial alpha of 0.05 based on the approximately 3,330 ‘blocks’ across the genome and our estimate of approximately 36 independent traits as determined by PCA on our phenotypes, resulting in a final genome-wide significance threshold of P=4.17E-7 and suggestive threshold of P=4.17E-6.

At our suggestive threshold we observe 72 loci across 25 distinct phenotypes for control CpG methylation affecting control traits, 39 loci across 16 phenotypes for isoproterenol-treated CpG methylation affected ISO-treated traits, 36 loci across 24 traits in which the *change* in CpG methylation was associated with a *change* in clinical traits and 32 loci across 19 phenotypes in which control CpG methylation levels were *predictive* of eventual ISO-treated clinical traits (All suggestive loci are detailed in Supplemental Table 7). At our genome wide significance threshold, we observe 12 loci across 8 phenotypes for untreated CpG and control phenotypes, 18 loci across 12 phenotypes for untreated CpG and ISO phenotypes, 25 loci across 12 phenotypes for treated CPG and ISO phenotypes and only 1 locus for delta CpGs and delta phenotypes (Table 1, Figure 3).

**Figure 3.**
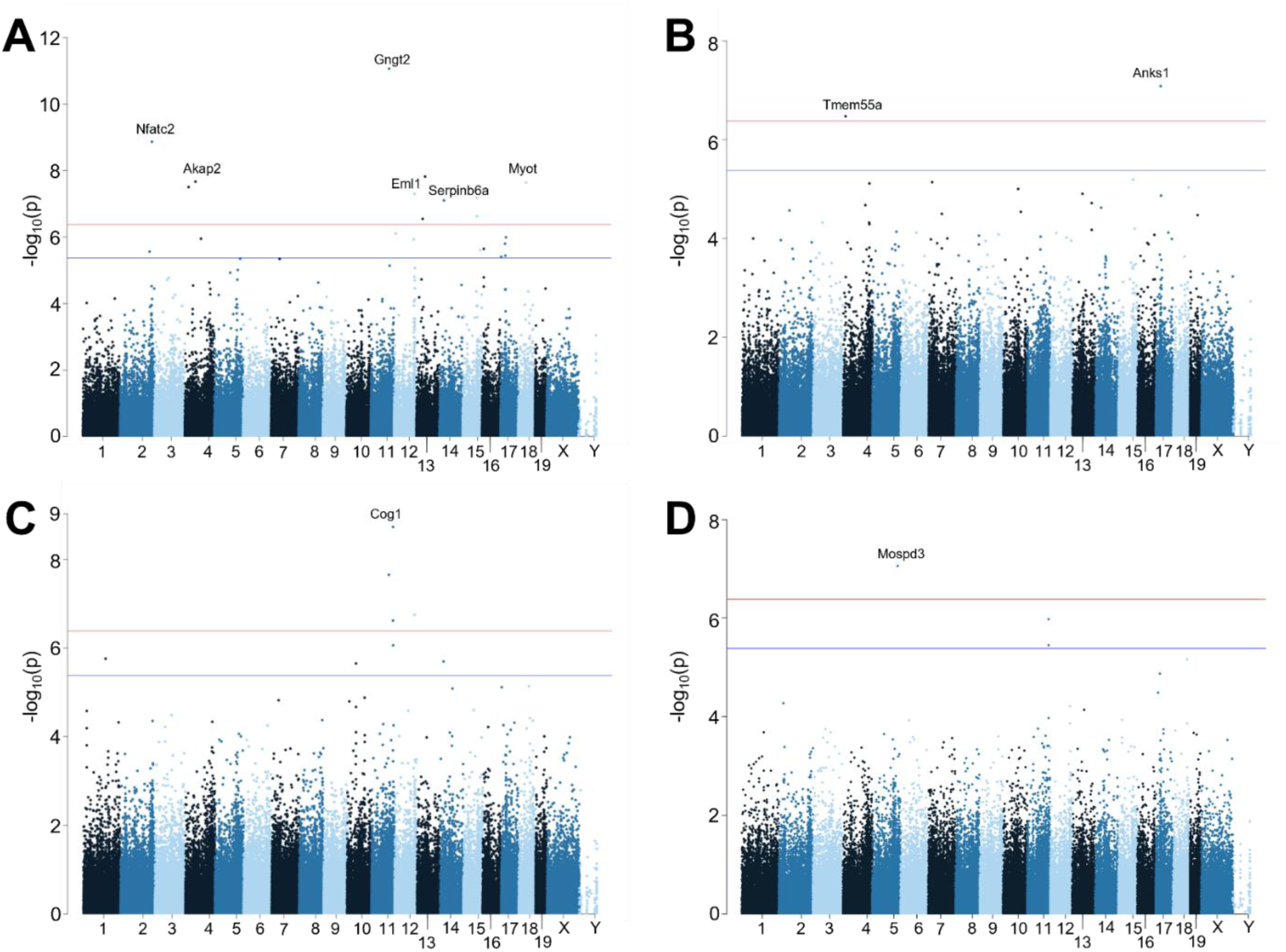
Representative Manhattan Plots from the EWAS Study. In each case, the X axis represents the position of a CpG across the genome and the Y axis is the negative log10 of the association p-value as determined by the MACAU algorithm. The red line indicates our calculated genome-wide significance threshold of P=4.17E-7, while the blue line denotes our suggestive threshold of P=4.17E-6. Genes of interest are highlighted and detailed further in Table 3. **A)** Treated CpGs affecting Isoproterenol Cardiac Fibrosis **B)** Treated CpGs affecting Isoproterenol Posterior Wall Thickness. **C)** Untreated CpGs affecting Control Adrenal Gland Weight **D)** Treated CpGs affecting Treated Adrenal Gland Weight.

**Table 1.**
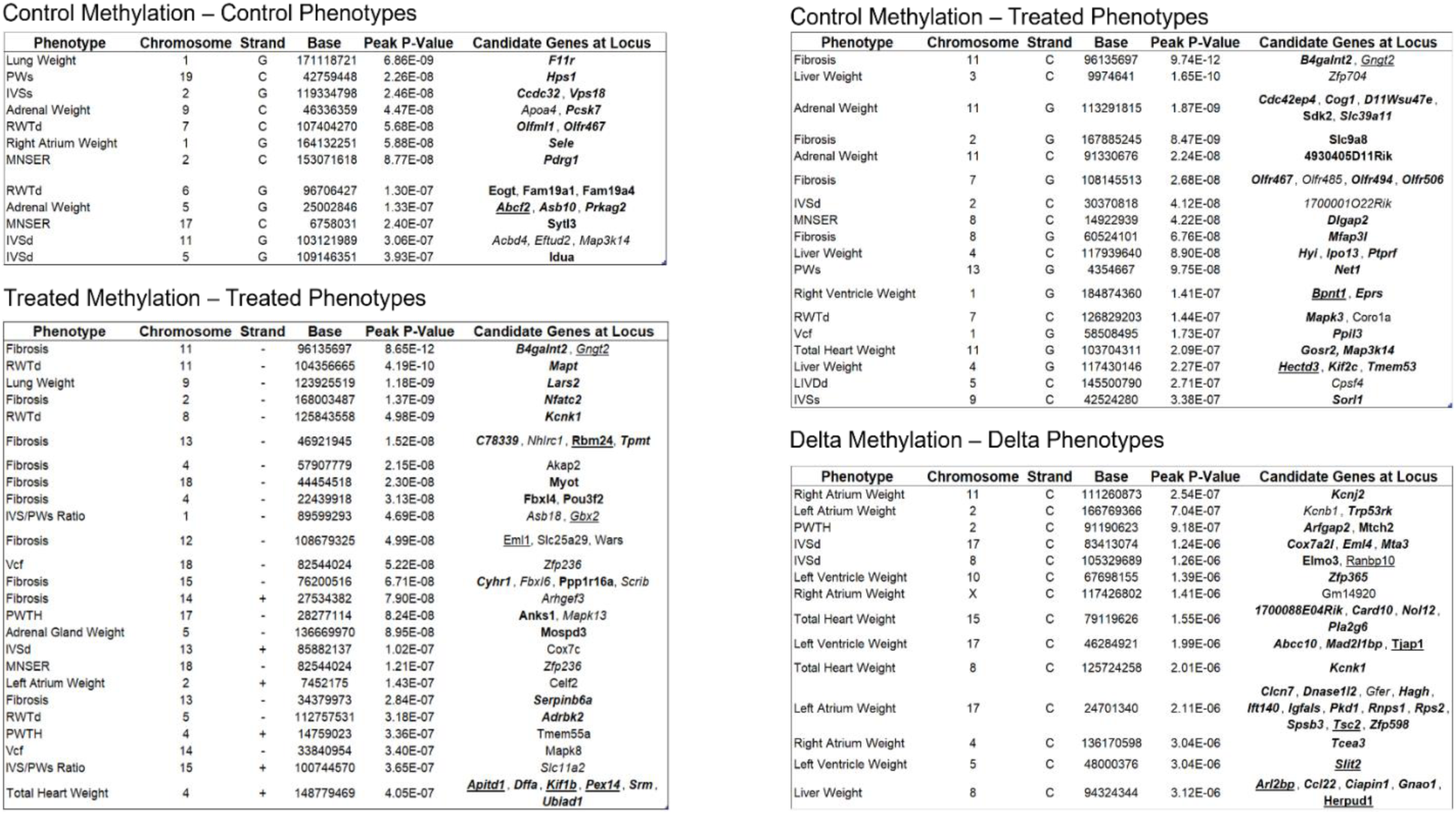
Methylome Loci in the HMDP Heart Failure Study. Genome-wide Significance Threshold – 4.18E-7, Suggestive Threshold 4.18E-6. IVS – Intraventricular Septum MNSER – Mean Normalized Systolic Ejection Rate PW -Posterior Wall PTH - Posterior Wall Thickness RWT – Relative Wall Thickness Vcf – Velocity of Centrifugal Force

### EWAS replicates previously identified GWAS loci and identifies novel associations

We have previously performed a GWAS for HF-associated phenotypes in this same panel of mice^5,6^. In prior studies in the HMDP, we have identified the average LD block size for the HMDP to have a resolution of approximately 2 Mb^33^. As such, to look for overlaps between GWAS and EWAS associations, we looked for any pair of GWAS/EWAS terms which lay within 2 Mb of one another. At our suggestive threshold for both GWAS and EWAS, we observe 209 EWAS loci and 41 GWAS loci for phenotypes analyzed by both approaches. Of these, 20 EWAS loci (9.6%) were within 2 Mb of a GWAS locus, suggesting possible co-regulation at that locus. This represents a modest, but significant enrichment over what would be expected by chance (P=0.0132).

One example of an overlapping EWAS/GWAS locus is found on chromosome 5 at approximately 136.7Mb. This locus is significantly associated with isoproterenol-treated RV weight by GWAS (P=3.49E-10)^5^ and isoproterenol-treated adrenal gland weight by EWAS on treated CpGs (P=8.95E-8). The best candidate gene at this locus is *Mospd3*. *Mospd3* is a poorly characterized gene that was first described in a manuscript that suggested that its knockout leads to a not fully penetrant thinning and occasional rupturing of the right ventricular cardiac wall during development^35^. Since 2020, additional reports have suggested that *Mospd3* may play a role in the regulation of mitochondria-ER binding and may help modulate mitochondrial membrane refreshment^36^.

Another example is *Prkag2*, also found on chromosome 5 at approximately 25.0Mb. Like *Mospd3*, this gene is also associated with both RV weight in GWAS (P=1.23E-6)^5^ and adrenal gland weight in EWAS (P=1.33E-7) after isoproterenol stimulation. *Prkag2* mutations cause an autosomal dominant glycogen storage disorder characterized by significant cardiac hypertrophy and subsequent heart failure^37^.

### EWAS loci contain known and novel candidate genes

In addition to the overlapping loci detailed above, we also identified a number of novel loci for this study (Figure 3, Table 1). To move from loci to candidate genes, we leverage the extensive ‘omics resources that our group has developed for the HMDP, including information at the genomic, transcriptomic, and phenotypic levels to identify and prioritize genes within our loci for downstream *in vitro* confirmation studies. We began by identifying all genes with 1 Mb upstream or downstream of the peak CpG in each EWAS locus. We next examined these genes to identify features that increase their likelihood of being causally involved with our phenotype, such as mis-sense or non-sense mutations as captured by the sequencing efforts of the Wellcome Trust Mouse Genomes Resource^28,29^, changes in gene expression associated to either SNP^5^ or methylation changes at the locus across the HMDP population, and whether prior literature supports the role of the gene in regulating changes in the phenotype and/or DNA methylation. Using these criteria, we were able to identify at least one gene per genome-wide significant CpG locus that showed sufficient evidence for further study, with many loci containing several genes implicated by multiple forms of evidence.

Contained within these loci are a number of genes which have already been associated with heart failure or other cardiomyopathies by other researchers. These include *Nfatc2*, the only candidate genes within a locus associated with cardiac fibrosis (P=1.4E-9) on chromosome 2 and reported to be a necessary mediator of calcineurin-dependent heart failure^39^. This connects with our prior research which linked multiple subunits of calcineurin to cardiac dysfunction in the HMDP^5^. We also observe *Celf2*, located on chromosome 2 and associated with atrial weight (P=1.4E-7). *Celf2*, also known as *Cugbp2*, works in opposition to *Celf1* to regulate mRNA stability and splicing^40^ and the *Celf* family has been implicated in multiple forms of cardiomyopathies and dysfunction^41,42^. Finally, knockout of our candidate gene, *Mapt,* at the most significant locus for Relative Wall Thickness after treatment (P=4.2E-10) has been shown to lead to diastolic heart failure^43^.

A number of loci contain genes which show clear involvement in the heart and make excellent candidates for further analysis. Several of these promising candidates are channel proteins, including our candidate gene in our single significant delta locus, where change in DNA methylation after ISO is associated with changes in phenotypic traits. *Kcnj2,* which is associated (P=2.5E-7) with changes in atrial weight, is a subunit of the sodium-potassium channel Kir2.1, and is the only known causal gene for Andersen-Tawil syndrome, which is characterized by ventricular arrythmias and other dysfunctions driven by an inability to properly process adrenergic stimuli^44^. We also observe *Akap2*, a gene which acts to slow deleterious cardiac remodeling by promoting angiogenesis and blocking apoptosis through the *Akap2/Pka/Src3* complex^45^ as well as regulating the migration of activated myofibroblasts in the establishment of cardiac fibrosis^46^ and which is associated in our data with cardiac fibrosis (P=2.2E-8). We further observe *Mapk8*, associated with Vcf (3.4E-7), that we previously showed was transcriptionally associated with right ventricular hypertrophy in a swine model of HFpEF^47^.

Still other genes represent novel targets with minimal evidence or associated mechanisms related to heart failure which our research highlights for potential downstream investigation. For the sake of brevity, we will only focus on a few interesting candidates. These include our best candidate for our most significant control-treated locus for fibrosis (8.6E-12) on chromosome 11, *Gngt2*. *Gngt2* is canonically a regulatory subunit of transducin, and was originally reported as playing a key role in phototransduction^48^. More recently, it has also been highlighted as a potential SNP for dilated cardiomyopathy in a Chinese population^49^ while its knockout in mice by the International Mouse Phenotyping Consortium (IMPC) leads to increased anterior wall thickness^50^. Similarly, knockout of *Anks1*, associated with posterior wall thickening (P=8.2E-8), is reported to lead to reduced posterior wall thickness in the IMPC^50^, but its role has never been reported on in the broader literature, although its family of Ankyrins has been implicated in cardiomyopathies more generally^51^.

### In vitro knockdown of candidate genes results in altered cellular dynamics in NRVMs

After identifying a number of promising candidate genes for cardiac phenotypes through ‘omics analyses of our identified loci, we sought to validate several of our candidate genes *in vitro* through siRNA-mediated knockdown of these candidate genes in Neonatal Rat Ventricular Cardiomyocytes (NRVMs).

As a proof of concept, we first targeted *Anks1*, whose knockout is associated with reduced wall thickness in the IMPC as discussed above, but whose role in the heart beyond this phenotyping report is unclear. We knocked out *Anks1* with a siRNA (IDTDNA, see Supplemental Table 2) in NRVMs, observing a ∼60% reduction in gene expression compared to scramble control. (Figure 4B). We are able to confirm the IMPC results, showing a 24% reduction in NVRM cross-sectional area (P=3.4E-8) at baseline and a 33% reduction after ISO treatment (P=9.3E-14) (Figure 4C). *Anks1* knockdown also blunted the effects of ISO, which increased *Anks1* KD NRVM cross-sectional areas by only 8% (P=0.06) whereas scramble+ISO cross-sectional areas increased 23% (P=2.7E-5).

**Figure 4.**
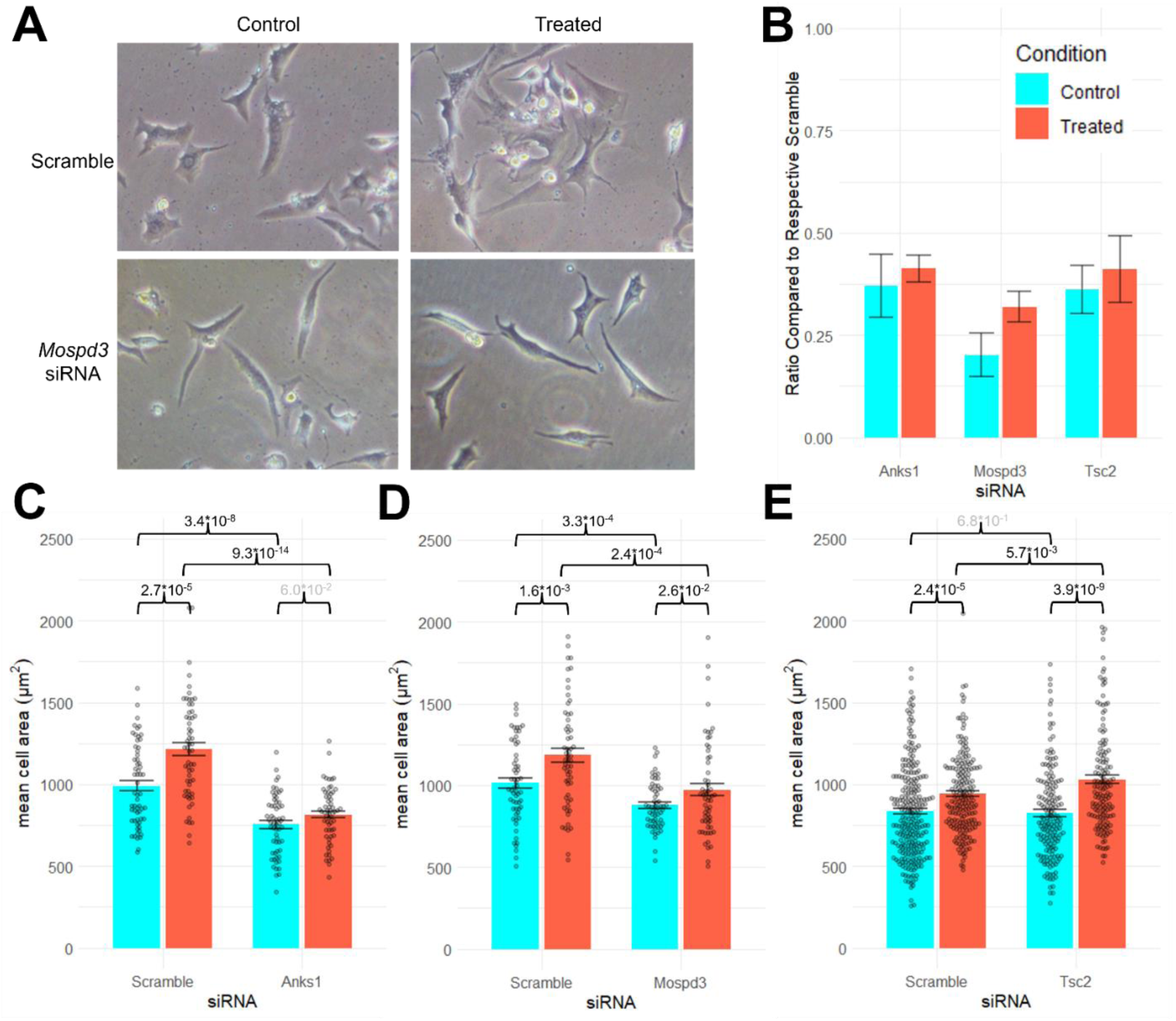
*in vitro* NRVM studies of candidate gene knockdown. **A)** Representative image of 20x resolution NRVMs **B)** percentage of siRNA-targeted gene expression compared to scramble controls. N=9, representing 3 independent trials with 3 technical replicates each **C-E)** NRVM cross-sectional areas for scramble and siRNA-treated cells in both control and ISO-treated (60 uM) conditions. P values are indicated, with greyed-out p values deemed not significant. N for *Anks1* and *Mospd3* studies was 60 cells per siRNA/condition combination. N for *Tsc2* was 200 for Scramble Control and Scramble ISO, 175 for *Tsc2* Control and *Tsc2* ISO.

Next, we examined *Mospd3*, described above as the candidate gene within a locus that was discovered twice – once for treated right ventricular weight in GWAS^5^, and again in this study through EWAS for treated methylation to treated adrenal weight (Table 1). Knockdown of Mospd3 via siRNA(IDTDNA, X, Supplemental Table 2) in NRVMs resulted in 80% and 68% knockdown in control and treated conditions, respectively (Figure 4B). We observe that *Mopsd3* knockdown results in 14.5% smaller cardiomyocyte cross-sectional areas at baseline compared to scramble controls (P=3.3E-4) and 18% smaller areas after ISO treatment (P=2.4E-4).

*Mospd3* knockdown also appears to diminish but not eliminate the effects of ISO (17 vs 11% increase, P=1.6E-3 to P=0.026) (Figure 4D).

The third *in vitro* result we highlight features *Tsc2*, which is a candidate for change in atrial weight after ISO treatment on chromosome 17 (P=2.11E-6). *Tsc2*, or Tuberous Sclerosis Complex 2, is associated with cardiac rhabdomyomas, benign tumors present in 0.02% of children^52^. Of children with a rhabdomyoma, approximately 80% of them will have either a mutation in *Tsc1* or *Tsc2*^52^. Although rhabdomyomas have been associated with heart failure^52^, the effects of *Tsc2* knockdown alone is less clear, with a single article suggesting a possible role in cardiac hypertrophy consistent with our GWAS locus^53^. Unlike *Anks1* or *Mospd3* knockdown, knockdown of *Tsc2* (∼61% in both control and treated conditions (Figure 4B)) did not result in any significant change in cell size in untreated cells compared to scramble (1.1% increase, P=0.68), but instead exacerbated the effect of ISO on cross-sectional area compared to scramble (21% increase with knockdown, P=3.9E-9 vs 11% increase without, P=2.5E-5, Figure 4E).

We further performed *in vitro* knockdown of two additional genes (Supplemental Figure 3) Knockdown of *Coro1a*, a gene associated with Relative Wall Thickness at diastole on chromosome 7 (P=1.44E-7) that acts as an actin regulator and may play a role in cell shape and adhesion^54^, was associated with an insignificant effect on cross-sectional area in control NRVMs (P=.16), but a significant blunting of the effect of ISO (19% smaller than scramble treated cells, P=2.9E-5). Also, *Slit2*, associated with change in LV weight after ISO on chromosome 5 (P=3.04E-6). *Slit2* is a cell migration gene with a known role in cardiac development^55^ We observe after *Slit2* knockdown a global reduction in NRVM cross-sectional area (10% in control, P=7.3E-6, 8% in ISO, P=4.3E-7), but no observed effect of gene knockdown on the efficacy of ISO (34% increase in scramble cells, 37% in *Slit2* KD cells).

### DNMT inhibitor reverses effects of hypermethylation on gene expression in a susceptible mouse strain

Finally, we investigated whether pharmacological inhibition of DNA methyltransferase using N-phthalyl-L-tryptophan (RG108), a non-nucleoside inhibitor of DNA methylation^56,57^ would alter the phenotypic and transcriptional response to ISO stimulation. We selected the BTBRT<+>tf/J (BTBRT) strain as our significant responder strain as it showed a 57% increase in heart weight and 30% increase in ejection fraction after 3 weeks of ISO stimulation, with C57BL/6J (B6) as our control with a 22% increase in heart weight and a 1.5% decrease in EF. We set up three experimental conditions: 1) Saline 2) ISO (30mg/kg/day) and 3) ISO (30mg/kg/day) + RG108 (12.5 mg/kg/day) administered through Alzet osmotic minipump for 21 days using the original experimental setup^5^. At day 21, we observe that BTBRT mice given only ISO once again showed a severe HF response compared to saline control which was significantly rescued by RG108 administration (Figure 5A+B). In contrast, B6 showed a more modest shift in LVIDd and %EF after ISO only and no significant effect at the phenotypic level caused by the addition of RG108 (Figure 5A+B). Intriguingly, global DNA methylation for both strains was reduced in ISO + RG108 vs ISO alone by a similar degree (Supplemental Table 8), suggesting, as detailed in our EWAS results above, the likelihood that important phenotype-methylation changes are locus specific as opposed to pan-genomic.

**Figure 5.**
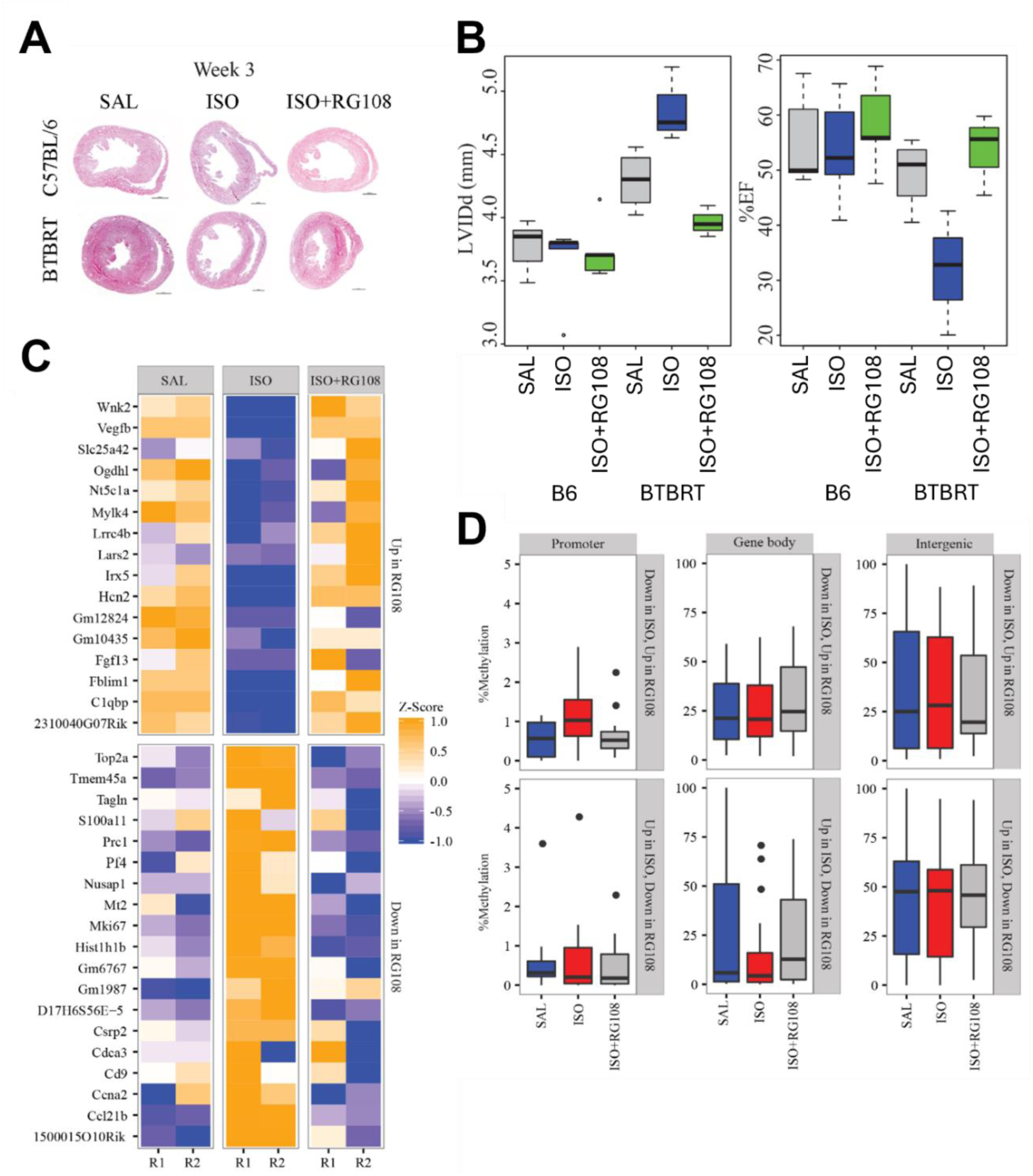
Concomitant DNMT inhibitor (RG108) treatment in ISO-treated mice de-methylates hypermethylated genes in severe-responder mouse strain BTBRT and is associated with an improvement in phenotype response. **A)** H&E cross-sections of hearts from “mild-responder” C57BL/6 and “severe-responder” BTBRT mice treated with SAL, ISO and ISO+RG108 for 3 weeks. **B)** Boxplot on C57BL/6 and BTBRT mice cardiac phenotype measurements LVIDd and %Ejection fraction after 3 weeks of RG108 treatment. **C)** Most significantly differentially expressed genes in BTRBT strain after RG108 treatment. Downregulated genes in ISO (blue) were upregulated in saline and ISO+RG108 (yellow). Similarly, the upregulated genes in ISO showed the opposite with downregulation in saline and ISO+RG108. D) Global DNA methylation distribution at the promoter, gene body and intergenic regions of the differentially expressed genes detected in Figure 3C. Corresponding images for B6 can be found in Supplemental Figure 4. N=12 per group.

To gain additional insights, we examined the effect of RG108 treatment on gene expression by performing differential expression (DE) analyses of RNAseq data gathered from ISO vs ISO+RG108 mice from both strains with the DESEQ R package^58^. In our significant responder strain, BTBRT, we observe 241 DE genes (q <0.05 & absolute LogFC >1.3) while in B6 we observe 327 DE genes at the same threshold (Supplemental Table 9, Figure 5C and Supplemental Figure 4). In both cases, most genes were upregulated after RG108 administration, although at a greater degree in in B6 (84% of all DE genes) compared to BTRBT (71%). 104 (43%) of the BTBRT DE genes are also observed in B6. Each of these genes show the same direction of fold change. Overlapping these DE genes with our EWAS hits revealed three EWAS candidate genes whose expression was affected by RG108 in both strains: *Mospd3*, which we describe above, along with *Lars2*, a tRNA synthetase with infrequent case reports suggesting a potential cardiac role^59^ and *Card10*, whose role in the heart is unclear but may be involved in pyroptosis^60^. No EWAS hits were unique to B6 mice, however we observed that *Akap2,* which we discuss above as a previously validated hit for regulating cardiac malformation^45,46^ was downregulated in BTBRT ISO vs Saline animals (log2FC -0.43), but restored in BTBRT ISO+RG108 mice (log2FC 1.75 vs ISO, 1.3 vs Saline). This suggests that *Akap2* may be a driving factor in the differential response to ISO in BTBRT compared to B6.

We sought to determine whether upregulation of these genes was the result of hypomethylation after RG108 treatment. We calculated the methylation levels of all the DE genes at their promoter, gene body, and intergenic regions across the three treatment conditions (Saline, ISO, ISO+RG108). At the promoter region, the downregulated genes in ISO were upregulated in RG108, displaying a contrasting distribution of increased methylation in ISO and a reduction in RG108 (Figure 5D). This finding is concordant with past studies where promoter methylation was found to be anti-correlated with gene expression^11,61,62^. In contrast, we observed minimal changes in DNA methylation at the genome body and intergenic regions across the three treatment conditions (Figure 5D).

## Discussion

In this study, we have performed a large-scale, genome-wide single-base resolution analysis of DNA methylation of hearts taken from 90 strains of the Hybrid Mouse Diversity Panel (HMDP), which consists of both classical inbred and recombinant inbred (RI) mouse lines under both control and isoproterenol-treated (30 g/kg/day for 21 days) conditions. Isoproterenol, a beta-adrenergic agonist administered through an implantable osmotic minipump in the abdominal cavity of these mice affords us a consistent means to induce cardiac hypertrophy and eventual heart failure while avoiding the effects of experimenter variability in models that rely on physical interventions such as coronary artery ligation to induce an infarction or trans-aortic constriction. In this study, we use tissue from the same animals we previously analyzed to study the role of DNA CpG methylation on hypertrophy and failure^5,6,16^.

To study the methylome of these animals, we performed reduced representational bisulfite sequencing (RRBS). RRBS is an affordable alternative to whole genome bisulfite sequencing (WGBS) that reduces the necessary number of reads per sample by limiting sequencing to regions of 200-500 basepairs flanked by *Msp1* digestion sites, enriching for CpG islands and sites near promoters and enhancers where DNA methylation is most likely to have an effect on gene expression and phenotypes^15,18^. We averaged 41.3 million uniquely aligned reads per sample across approximately 2.8 million CpGs. Filtering for CpGs present in at least 70% of the strains and at at least 5x coverage left us with 1.8 million CpGs, or approximately 8.4% of all CpGs in the mouse genome. This contrasts with a RRBS study performed in the livers of a different set of HMDP mice^25^ in which, despite reporting similar numbers for total aligned reads per sample (41.3 vs 41.0 Million), the prior study was able to capture 2 million CpGs at 10x coverage in at least 90% of the samples, a recovery rate of 9.6% despite a more stringent cutoff for inclusion. This relaxation of stringency is due to the increase in the number of samples (174 in our study vs 90 in theirs). As each RRBS outputs only a representative *sampling* of CpG sites rather than the full complement of sites which would be observed with WGBS or through the human-only Illumina Infinium methylome platform, increasing the number of samples *by necessity* decreases the number of CpGs which will reach a given coverage threshold.

Although this reduction in stringency does represent a limitation of our approach in that low-coverage CpGs have greater uncertainty compared to high-coverage CpGs, we were still able to identify a number of interesting candidates. In the future, deeper sequencing of these libraries may allow us to improve the rigor of our results.

For our first analysis, we limited ourselves to 168,251 ‘hypervariable’ CpGs – sites which differed by at least 25% absolute methylation in at least 5% of samples. We observe that only 397 (0.2%) of these hypervariable SNPs show a universal shift of at least 3% between control and treated mice at an FDR of 1% (Figure 1B). While the genes proximate to these sites are enriched for GO terms pertaining to apoptosis (P=1.8E-6) and abnormal cardiac morphology (P=2.7E-5) (Figure 1C), it is striking that so few CpGs show a universal response across all of our tested strains, suggesting that genetics rather than environment is a major driving factor of DNA methylation, at least in the context of our mice, whose differences in environmental exposure is limited to the presence or absence of ISO. Further supporting this is our analysis of the top 1,683 (1%) of varying CpGs across the HMDP regardless of the effects of ISO, where we observe that genetics, as shown by the RI panels and known related strains separating into distinct branches after hierarchical clustering (Figure 2, bottom edge) is much more apparent than the effects of ISO, which are not responsible for any sub-branch of the tree (Figure 2, top edge).

Motivated by our confirmation that genetic background plays a strong role in determining DNA methylation shifts, we queried whether these shifts were linked to cardiac phenotypes through an Epigenome-wide Association Study (EWAS) using the binomial mixed model approach MACAU which was specifically designed to work with RRBS count data^15^. We observe (Figure 3, Table 1) 56 significant loci in our study – 12 loci where untreated DNA methylation is linked to untreated phenotypes, 25 loci where treated methylation is linked to treated phenotypes, a single significant locus where the change in methylation is predictive of a change in phenotype, and, of greatest interest to us, 18 loci where untreated methylation levels were predictive of treated phenotypes. These predictive loci are a unique feature of EWAS when compared to GWAS studies. As DNA methylation can shift in response to environmental stimuli, being able to identify methylation states *before* environmental challenges that can then predict phenotypic responses *after* that challenge is a powerful tool for understanding potential mechanisms for the candidate genes identified in the more predictive (untreated CpGs to treated phenotypes) and more reactive (treated CpGs to treated phenotypes) loci.

In contrast to our GWAS hits^5,6^ in which we reported several loci that associated with the change of phenotypes after ISO stimulation, we observe only a single significant locus that links a change in methylation to a change in phenotype (Table 1). We view this as likely due to the increased levels of uncertainty in our measurements, where not only do we observe variability and noise in our phenotypic data at both control and treated conditions, but also in our methylation percentages. This significantly reduces the power we have to observe these sorts of loci. Additional strains of mice, characterization at the CpG and phenotypic level of additional mice per strain, and/or more precise means to measure DNA methylation may help to increase the number of change loci which researchers are able to recover.

We examined whether we observed GWAS/EWAS co-localization in our study, comparing the suggestive GWAS hits in our original studies^5,6^ to the suggestive EWAS hits from this study. We observe only a 9.6% overlap between our EWAS and GWAS loci, a result that, while technically significant (P=0.014), does not represent a broad consensus between our EWAS and GWAS hits. 9.6% is similar to the approximately 15% of EWAS/GWAS co-localizations that were observed in a prior HMDP EWAS/GWAS study in the liver{Orozco, 2015 #29;Bennett, 2010 #178}. This low overlap is likely due to a lack of statistical power in either our GWAS and/or EWAS studies to detect associations with small effect sizes. Orozco et al^25^ was able to show with gene expression EWAS and GWAS that molecular traits, whose regulation is significantly simpler than clinical traits, had a much larger overlap (77%) compared to their reported 15% for clinical traits. Of the sites that do overlap in our study, we observe a number of highly relevant candidate genes, such as *Prkag2*, or the gamma-2 subunit of the AMPK complex. Associated with right ventricular weight in the GWAS and adrenal gland weight in the EWAS after ISO stimulation, *Prkag2* mutations are known to be causal for an autosomal dominant form of cardiac hypertrophy^37,63^. Although we do not observe any evidence of full knockout of *Prkag2* in our cohort, our results do suggest that natural variation in *Prkag2* levels may be predictive of cardiac maladaptation to stressors independent of its KO-associated phenotype.

Beyond these overlapping loci, we also identified a number of loci which were unique to our EWAS study of heart failure. In many studies, moving from an identified locus to a list of likely candidate genes within that locus can prove challenging. In our study, however, we were broadly successful at identifying interesting candidates due to both the smaller ‘linkage’ blocks of correlated CpGs compared to SNPs (approximately 750kb in width compared to 2mb)^25,33^, the short range-of-action proposed for most CpGs^64^, as well as our ability to layer on additional forms of ‘omics data taken from the same mice that included detailed transcriptomics as well as sequencing data for each of the founder strains of the RI panels as well as other classically inbred lines^28^. Layering these data sources on top of one another highlights a few genes per locus as needing additional scrutiny (Table 3), greatly assisting in the identification of candidate genes within each locus. Several of the genes we flag within our loci have strong previous associations with cardiomyopathies, such as *Prkag2, Nfatc2, Akap2, Celf2* and *Mapt.* The presence of these genes increases confidence in our results. Our loci also contain candidate genes whose links to hypertrophy and heart failure are more tenuous, such as *Mospd3, Gngt2,* or *Anks1* and which deserve further scrutiny based on our findings.

We used primary neonatal rat ventricular cardiomyocytes (NRVMs) and siRNA-mediated gene knockdowns to study the role of several of our candidate genes *in vitro*. In some cases, we were able to replicate prior reported knockout or knockdown phenotypes. For example, we were able to show that *Anks1* knockdown reduced cardiomyocyte size in a manner similar to the reduced vessel wall thickness reported by the IMPC^50^, and extend these results by showing that *Anks1* knockdown also significantly blunted the effects of catecholamine stimulation in addition to its effects at baseline. Likewise, we validated the vessel wall thinning phenotype which is one of the only known features of *Mopsd3* knockout^35^, while suggesting a role for the gene in the regulation of heart failure beyond its previously reported role in heart development^35,36^. In other cases, such as with *Tsc2*, classically associated with cardiac rhabdomyomas^53^, we were able to show that gene knockdown was specifically associated with blunting the effects of ISO on cell size without affecting baseline cell size in untreated cells, suggesting a new avenue of functionality for this gene in the regulation of catecholamine-driven hypertrophy.

Finally, we asked what the effect of blocking the action of DNA methyltransferases (DNMTs) before catecholamine challenge using the methyltransferase inhibitor N-phthalyl-L-tryptophan (RG108) would have on cardiac phenotypes. We observe in our paired model of a severe responder to ISO challenge (BTBRT) and a more resistant strain (B6). We observe that DNMT knockdown in BTBRT was able to limit the effects of catecholamine-induced stress on the heart, maintaining Ejection Fraction and preventing chamber dilation, while the effects of RG108 on the resistant strain, B6, were not significant. As DNA methylation acts through the regulation of genes to affect phenotype, we next focused on the differentially methylated genes in both strains, observing similar numbers of DE genes in each with a 43% overlap. GO enrichment of these genes highlighted enrichment for cardiac contraction genes (Supplemental Table 10) and analysis of changes in promoter methylation status of the DE genes in the BTBRT strain showed that RG108 prevented the hypermethylation seen in ISO animals. Overlapping these DE genes with our EWAS hits highlighted three genes in common to both B6 and BTBRT, namely *Lars2, Card10* and *Mospd3*, further highlighting the latter’s need for further study, while *Akap2*, a gene that acts to reduce cardiac remodeling through control of angiogenesis and apoptosis through the *Akap2/Pka/Src3* complex^45,46^, is downregulated in BRTBT ISO vs BRTBT saline (log2 fold change -0.43) and restored in BRTBT ISO+RG108 (log2 fold change 1.75 vs ISO, 1.3 vs saline), suggesting that changes in *Akap2* methylation and subsequent gene expression changes may be directly related to BTBRT’s more significant response to ISO stimulation.

Our study has some limitations. Firstly, our use of only female mice hinders our ability to easily extend our findings to male mice. Due to cost constraints, during our pilot study we observed that there was a greater variation of response to ISO in female mice among the parental lines of the RI panel that makes up the majority of the HMDP (A/J, C57BL/6J, C3H/HeJ, DBA/2J) and chose to maximize our ability to recover loci of interest by focusing only on female mice. While this is a limitation, past studies in the HMDP^65,66^ suggest that many loci identified in female mice are also observable in male mice. A second limitation concerns the variability of cell type proportions within the mammalian heart and its effects on DNA methylation. Multiple reviews^67–69^ have highlighted the difficulty in accurately measuring the proportion of cell types (e.g. Cardiomyocytes, Fibroblasts, Endothelial Cells, etc.) within the heart, with significantly different results based on species, location within the heart, method of study, individual analyzed, etc. There is no good understanding of the variability of cell type proportions in the heart within a population, for example. Additionally, disease processes shift these proportions in unequal ways depending on genetic background. For example, we observed in our GWAS that mutations in the *Abcc6* gene led to significant apoptosis of cardiomyocytes and replacement with fibrotic tissue^5^. Differences in DNA methylation are one of the major ways in which cell types are differentiated from one another^8,25,70,71^. Shifts in cell-type proportion are a known and appreciated confounder to EWAS approaches^34^, typically addressed by introducing covariates that account for the relative proportions of cell types to one another across the cohort. While this is easily achieved in some tissues (e.g. blood), it has proven very difficult to ascertain in the heart, and likely affects our identified loci through both amplifying the signal of loci associated with genes involved in specific cell types, while suppressing other signals. We feel this is one of the major reasons why many of the most significant loci we recovered were for cardiac fibrosis, whose link to increased or decreased relative numbers of fibroblasts is clear. Finally, the use of RRBS instead of WGBS likely led to sampling error and reduced power which could be counteracted through an increased depth of sequencing or the addition of more strains.

Cardiac hypertrophy and remodeling are major determinants of HF progression. Our results represent new avenues of investigation into the genomic locations and gene transcripts which drive these phenotypes. Our use of cardiac tissue and careful high-throughput integration of molecular phenotypes such as cardiac transcriptome and methylome data has highlighted a number of interesting and novel candidate genes and represents a powerful alternative to human studies which are frequently limited in terms of sample size, environmental noise, and multi-omic integration. Further refinement of our loci and the addition of additional data such as cardiac cell composition will further shed light on the role of the methylome in the progression of heart failure with the ultimate goal of improved personal therapies for patients.

## Supporting information

All Supplemental Tables

## Data Availability

RRBS data from the HMDP is available at the Sequence Read Archive at accession PRJNA947937. RRBS data from the RG108 experiments is available at the Sequence Read Archive at accession PRJNA945923. Gene Expression from the HMDP is available at the Gene Expression Omnibus at accession GSE48760. HMDP Phenotypic data is available through Mendeley Data at accession 10.17632/y8tdm4s7nh.1.

## Acknowledgements

CDR, CL, AD were supported by R00HL138301 and R01HL162636. We thank Douglas Chapski, PhD for his help with the methylome processing pipeline.

