## Supplemental Figures 1,3,4 for "Mapping DNA Methylation to Cardiac Pathologies Induced by Beta-Adrenergic Stimulation in a Large Panel of Mice"

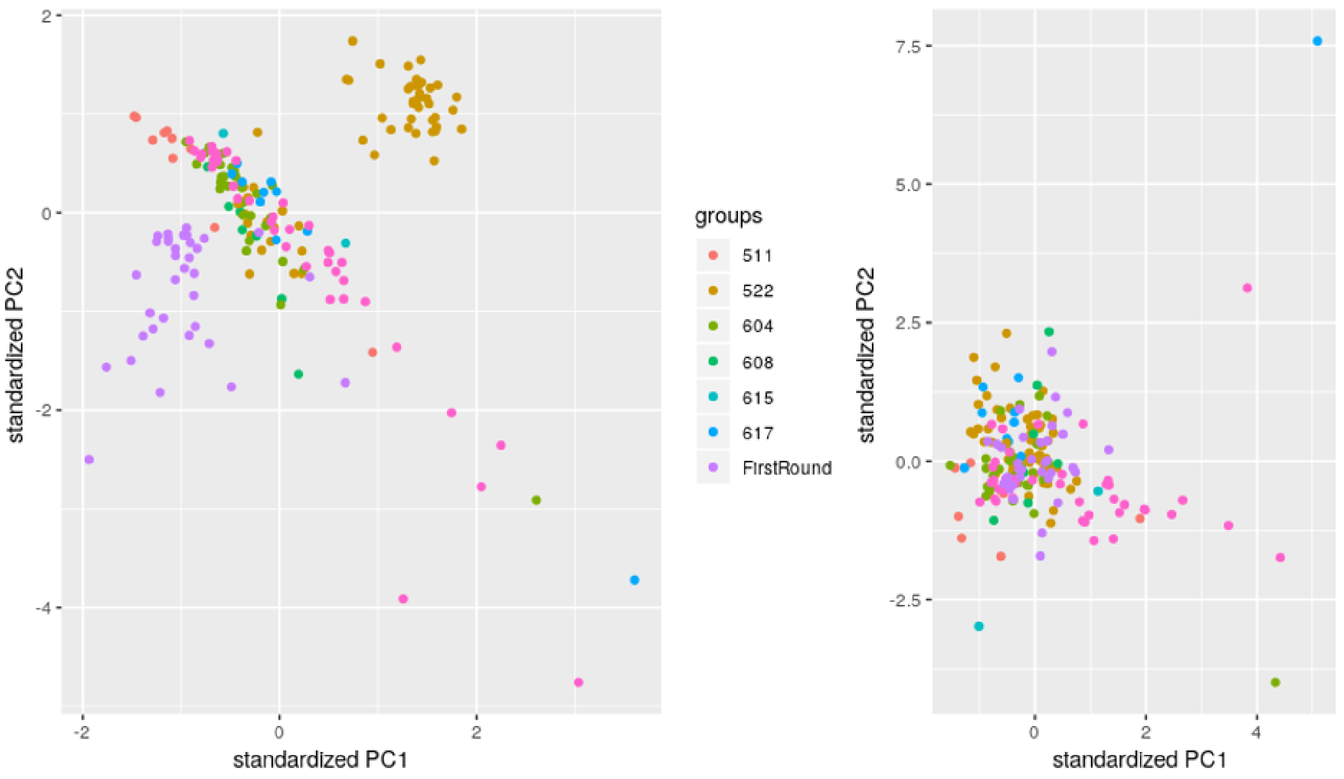


Supplemental Figure 1 - **Batch Effects pre and post correction via COMBAT.** PCA of all samples before (left) and after (right) correction. On the left, the batch effects due to date run (511 = 5/11, 604 = 6/4, etc) are plainly obvious. On the right, these batches have been largely eliminated.

Supplemental Figure 2 - A larger version of the heatmap from Figure 2A with gene names clearly visible.

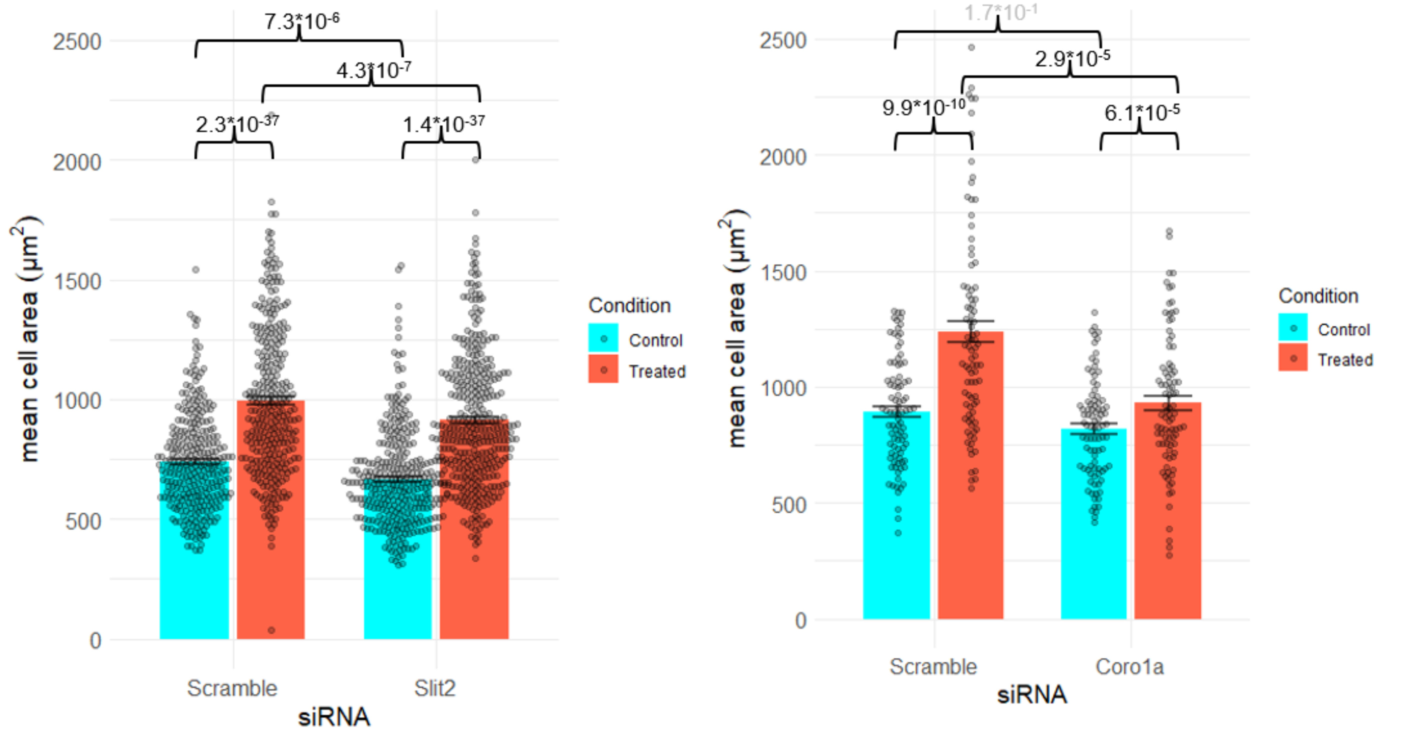


Supplemental Figure 3 - **SiRNA-mediated Knockdown of Slit2 and Coro1a in NRVMs.** Effect of gene knockdown with either scramble siRNA or *Slit2* or *Coro1a* siRNAs in both control and ISO-treated conditions. Numbers indicate p-values via Student's t-test. Greyed-out numbers are not significant. Slit2 N=225 per condition . Coro1a N= 60 per condition.

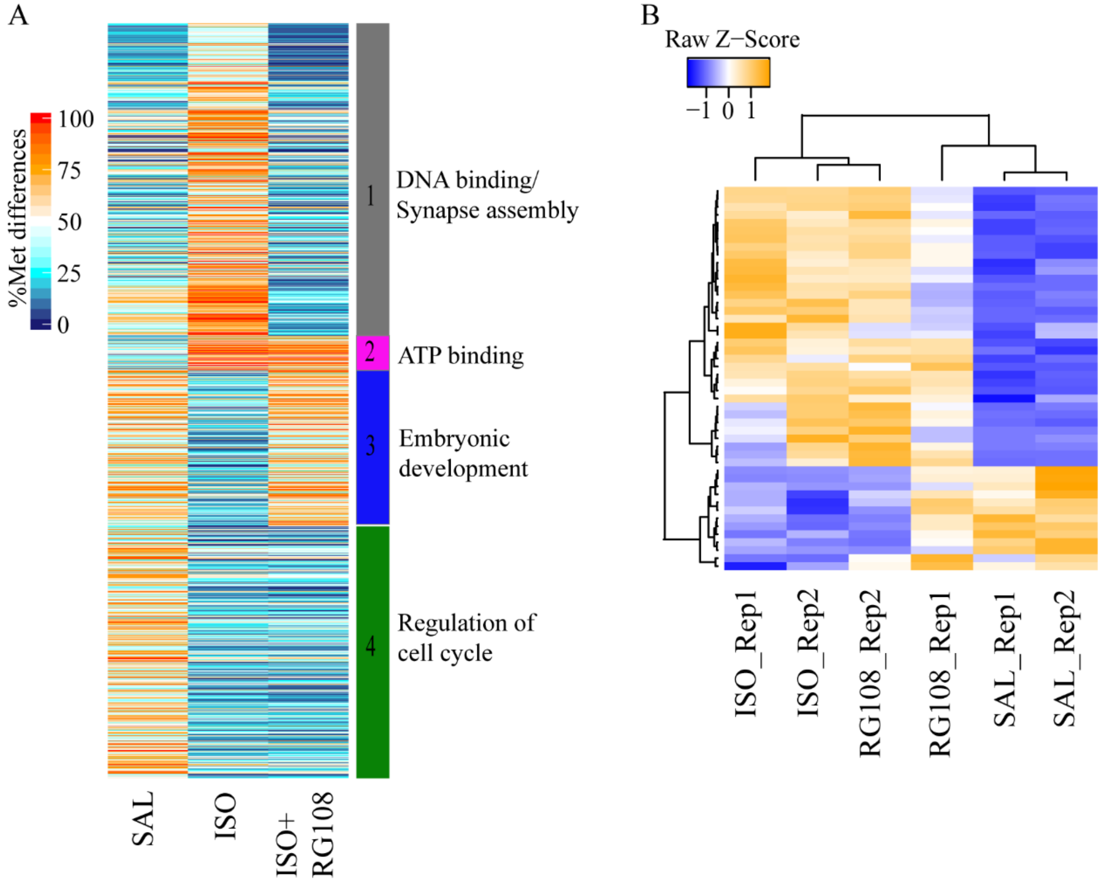


**Supplemental Figure 4. Differentially methylated regions and gene expression in C57BL/6.** A) Differentially methylated regions (DMRs) (q-value <0.01, PM >25%) in C57BL/6 identified through pairwise comparison across SAL vs ISO, SAL vs ISO+RG108 and ISO vs ISO+RG108. DMRs identified in each pairwise comparison were further cross compare to determine which groups of DMRs show consistent methylation changes between each treatment conditions. 4 distinct groups were identified (1) DMRs reversed after ISO+RG108 treatment (2) DMRs hypermethylated in both ISO and ISO+RG108 (3) DMRs hypermethylated after RG108 and (4) DMRs which remained hypomethylated in both ISO and ISO+RG108 treated. B) Differential gene expression (q-value <0.05 and log2FC >1.3) across SAL vs ISO, SAL vs ISO+RG108 and ISO vs ISO+RG108. No major differential expression differences was observed between ISO and ISO+RG108.
