## Supplementary figures and images for "Mapping DNA Methylation to Cardiac Pathologies Induced by Beta-Adrenergic Stimulation in a Large Panel of Mice"

### Supplemental Figure 2.png

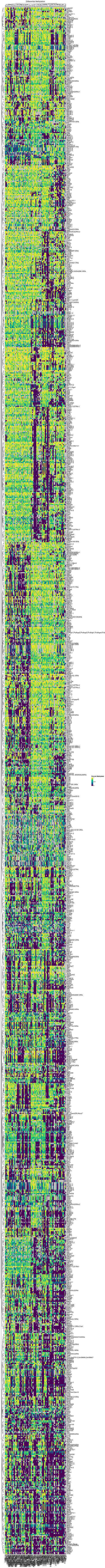
